# Temperature-mediated dynamics: unraveling the impact of temperature on cuticular hydrocarbon profiles, mating behavior, and life history traits in three *Drosophila* species

**DOI:** 10.1101/2024.01.20.576429

**Authors:** Steve B. S. Baleba, Nan-Ji Jiang, Bill S. Hansson

## Abstract

In a world grappling with climate change, understanding the enduring impact of changes in temperature on insect adult traits is crucial. Here, we explored the intricate dynamics of exposure to different temperatures in three *Drosophila* species: *Drosophila ezoana* originating in Arctic regions, *D. novamexicana* in arid, hot environments, and in the cosmopolitan species *D. virilis*. Rearing these flies at 15, 20, 25, and 30°C revealed striking variations in their cuticular hydrocarbon (CHC) profiles, known to mediate mate recognition and prevent water loss in insects. The cold-adapted *D. ezoana* consistently exhibited reduced CHC levels with increasing temperatures, while the warm-adapted *D. novamexicana* and the cosmopolitan *D. virilis* displayed more nuanced responses. Additionally, we observed a significant influence of rearing temperature on the mating behavior of these flies, where those reared at the extreme tempreatures, 15 and 30°C, exhibiting reduced mating success. Consequently, this led to a decrease in the production of adult offspring. Also these adult offspring underwent notable alterations in life history traits, reaching adulthood more rapidly at 25 and 30°C but with lower weight and reduced longevity. Furthermore, among these offspring, those produced by the cold-adapted *D. ezoana* were more vulnerable to desiccation and starvation compared to those from the warm-adapted *D. novamexicana* and the cosmopolitan *D. virilis*. In summary, our research underscores the intricate interplay between temperature, ecological adaptation and various life history traits in *Drosophila* species from distinct agro-ecological regions. The observed variations in CHC profiles, mating behavior, fertility and responses to environmental stressors collectively provide valuable insights into how environmental conditions shape the biology and ecology of insects.

## Introduction

Butterflies, bees, moths, beetles, and flies all display a complete metamorphosis (holometaboly) as they progress through their developmental stages before reaching adulthood (Rolff et al., 2019). In these insects, the immature stages (eggs, larvae and pupae) have no physical resemblance to the adult stages and undergo substantial morphological and physiological transformations to complete their development. In this process, the hatching egg produces a larva that undergoes multiple moults (or ecdysis), during which the old cuticle is shed and replaced with a new one. This transformation ultimately leads to the formation of a pupa, from which an adult will eventually emerge (Hall and Martín-Vega, 2019). During this transformation, a significant restructuring occurs in larval tissues and organs, involving the nervous system, midgut, muscles and cuticle (Tettamanti and Casartelli, 2019). Going these changes can impose considerable stress and challenges for the insects. Any environmental conditions that are inconsistent or unfavorable can further intensify this stress, potentially impeding the success of the transformation process. A varying environment often triggers plasticity that generates multiple phenotypes in insects with the same genome (Yoon et al., 2023).

Among insect organs, the cuticle is highly susceptible to environmental changes due to its direct exposure to the external world. Within this organ, specialized cells called oenocytes secrete cuticular hydrocarbons (Schal et al., 1998), which often serve a dual role (Chung and Carroll, 2015; Holze et al., 2021). They protect terrestrial insects against desiccation. For example, using 50 drosophilids, Wang et al. (2022) demonstrated that the composition of CHCs can explain up to 85.5% of the variability in desiccation resistance. Also, CHCs serve as signaling molecules in a wide variety of chemical communication systems. They serve as the primary signals to attract conspecific mates for courtship and copulation behaviors, as well as to indicate receptivity, fertility, and mating status (Kabir et al., 2023; Khallaf et al., 2021). The production of cuticular hydrocarbons in adult insects can be influenced by the stress experienced during preimaginal development. For example, in *Drosophila mojavensis*, adults reared on their natural host (cactus) produced more CHCs as opposed to those reared on laboratory food (Etges and Oliveira, 2014). Eggs of *D. melanogaster* (Rajpurohit et al., 2017a, 2021; Savarit and Ferveur, 2002) and *D. mojavensis* (Gibbs et al., 1998) incubated under different temperature conditions give rise to adults with different CHC profiles. However, the extent to which these changes in CHC profiles can impact the mating behavior of insects and the fitness of their subsequent offspring remains uncertain.

The current climate changes, primarily driven by human activities, are causing a rise in global temperatures (Kabir et al., 2023; Winkler et al., 2020). As a consequence, insects are encountering temperatures beyond what has been present in their historical ranges (Deutsch et al., 2008; Ma et al., 2021). It is postulated that holometabolous insects may exhibit variable thermal adaptations and responses to suboptimal temperatures based on their geographical distribution (Hoffmann, 2010; Kingsolver et al., 2011; Porcelli et al., 2017). As reviewed by Hodkinson (2005), this adaptation to local climate conditions can happen through shifts in insect behavior (e.g. mating frequency and duration), fertility, life history traits (e.g. developmental time, body size and longevity) and physiological attributes (e.g. desiccation and starvation resistance). Here, we aimed to investigate how rising rearing temperatures affect the reproduction, fitness, and physiology of insects living in different latitudinal zones. We hypothesized that the composition of the CHC layer in drosophilid species from different latitudes would change with rising temperatures. This in turn would have a significant impact on their mating behavior, fertility, on different life history traits (ie. developmental time, body size, longevity), as well as on desiccation and starvation resistance of their subsequent offspring. To test our hypothesis, we selected three closely related species within the virilis group: *Drosophila ezoana*, *D. novamexicana* and *D. virilis*. These species have adapted to inhabit subarctic, desert, and global climates, respectively, and display distinct temperature preferences (Baleba et al., 2023). They all feed and breed on slime flux, saps, and decaying bark of tree species, predominantly from the Salicaceae and Betulaceae families (Throckmorton, 1982).

We analyzed the cuticular hydrocarbon compositions of adults of the three drosophilid species when developed at 15, 20, 25, and 30°C. Next, we investigated whether these flies exhibit distinct mating behavior depending on reaing temperature. Also, we assessed their fertility and determined life history traits of their offspring. Finally, we tested how well the offspring could endure extended periods of extreme dryness (desiccation) and food deprivation (starvation). Our study provides comprehensive insights into how insects from various latitudinal regions are likely to respond to the ongoing global warming.

## Material and methods

### Fly stocks and maintenance

We obtained *Drosophila ezoana* (stock number: E-15701) from the fly stocks of Kyorin university, Japan (https://shigen.nig.ac.jp/fly/kyorin/index_ja.html), while *D. novamexicana* (stock number: 15010-1031.08) and *D. virilis* (stock number: 15010-1051.00) were obtained from the National *Drosophila* Species Stock Center of Cornell University (https://www.drosophilaspecies.com/). We reared *D. ezoana* at 20°C, 16h Light: 8h Dark, *D. virilis* at 23°C 12h Light: 12h Dark, and *D. novamexicana* at 25° C 12h Light: 12h Dark. All flies were fed on autoclaved cornmeal-yeast-sucrose-agar food.

### Flies preparation and cuticular hydrocarbon analysis

Gravid females of *D. ezoana*, *D. novamexicana*, and *D. virilis* from the stocks were allowed to lay eggs in food vials at low density (20-30) for 5 hours. These vials were immediately transferred inside four Snider Scientific incubators (www.snijderslabs.com) set respectively at 15, 20, 25 and 30°C, all with 70% humidity and 12h Light:12h Dark. Once emerged, flies were collected within 1 hour, sexed and returned to their respective developmental incubator. 7 days later, 5 flies of the same sex from each incubator were soaked for 10 minutes in hexane (Supplementary Fig. 1). containing 1-Bromoeicosane (25 ng/μl) as an internal standard (Savage et al., 2021; Wang et al., 2022). Each body extract sample was analyzed by gas chromatography coupled with mass spectrophotometry (7890B GC System, 5977A MSD, Agilent Technologies,https://www.agilent.com) equipped with a polar column (HP-INNOWAX, 30 m long, 0.25 mm inner diameter, 25 μm film thickness; Agilent) with helium as carrier gas. Each sample (1 μL) was injected into the GC-MS with an autosampler (Agilent Technologies). The inlet temperature was set to 250° C. The temperature of the GC oven was held at 50°C for 2 min, increased gradually (15°C/min) to 250°C, which was held for 3 min, and then to 280°C (20°C/min) and held for 30 min. The MS transfer line, source, and quad were held at 280, 230, and 150°C, respectively. GC-MS data were processed with the MDS-ChemStation Enhanced Data Analysis software (Agilent). For each developmental temperature and species, we injected 5 samples per sex. In every sample, we determined the quantity of each identified CHC by comparing its peak area with that of the internal standard (normalized intensity).

### Mating assay

To investigate the impact of temperature on the mating behavior of *D. ezoana*, *D. novamexicana*, and *D. virilis*, virgin individuals of these flies reared at 15, 20, 25, and 30°C were separated by sex upon emergence, placed in groups (20 individuals per vial), and returned to their respective incubator. Seven days later, a single male and a single female from each temperature condition were introduced (using a mouth aspirator) in a courtship chamber and their behavior was observed for 1 h (Supplementary Fig. 1). Each courtship arena was equipped with 4 chambers (1 cm diameter × 0.5 cm depth) covered with a plastic slide. A constant airflow of 0.2 mL/min was added from below to each arena. To record the courtship behaviors of the couples, the four chambers were observed simultaneously using a GoPro Camera 4 or Logitech C615. For each developmental temperature, we observed 48 couples. To quantify the mating behavior of each pair, we manually examined each video to determine mating success (i.e., the percentage of males that managed to mount the females) and mating duration (i.e., the amount of time a male spent on top of the female). All these mating assays were performed at 23°C and 70% humidity.

### Fertility assay

To elucidate how temperature impacts the fertility of *D. ezoana*, *D. novamexicana*, and *D. virilis*, each mating pair that managed to mate in the previous experiment was gently collected (using a mouth aspirator) and placed in a vial containing fresh food (13 g) and transferred to their corresponding incubator (previously set at 15, 20, 25, or 30°C). For each couple, we allowed the female to lay eggs throughout its lifespan and closely observed the progress of egg development until the emergence of adult offspring. This allowed us to count the total number of adult offspring (fecundity) produced by each species at the different developmental temperatures. Each mating pair was transferred to a new food vial each day.

### Determination of life history traits

During the fertility test, we simultaneously collected data on developmental duration (from egg to adult), pupal weight, and adult body weight (both fresh and dry). For each experimental temperature, we weighed 30 pupae/species individually using a Sartorius balance (0.0001 g precision). To determine the fresh adult body weight of each species, we placed 10 males and 10 females from each temperature group into pre-weighed 1.5 ml Eppendorf tubes and recorded the total weight using a Sartorius balance. For each developmental temperature, species, and sex, 10 independent trials were conducted. To measure the dry body weight in each trial, we took these tubes, placed them in an oven set at 40°C for 10 minutes to kill the flies, and then transferred them to room temperature. We subsequently weighed them every two days until a constant weight was noted. The fresh and dry body weigh was obtained by subtracting the weight of the Eppendorf tube from the weight of the tube containing the fresh and dry flies, respectively. Under each developmental temperature (15, 20, 25, and 30°C), we also tested the longevity of the adult offspring produced by each species. To do this, we monitored weekly (until the death of the final individual), newly emerged adult offspring (20 individuals/sex/food vial). In each trial, flies were transferred into a new food vial every week and we used 10 replicates.

### Desiccation and starvation resistance assays

In addition to the precedent life history traits, we also examined the survival capacities of the adult offspring when confronted with prolonged periods of severe dryness (desiccation) and food deprivation (starvation). We evaluated desiccation resistance using the methodology outlined by Chung and Carroll (2015). For each developmental temperature and species, we exposed 10 adult offspring (7 days old) of the same sex to an assay setup (see Wang et al. (2022)) containing 10 g of silica gel (S7500-1KG, Sigma-Aldrich, Darmstadt, Germany). We recorded mortality every 2 hours until the last individual succumbed. For the starvation resistance assays, we followed the protocol described by Nayak and Mishra (2019). Here, for each developmental temperature and species, 10 adult offspring (7 days old) of the same sex were removed from the food vial and transferred to a tube containing a moistened cotton plug at the bottom (to prevent death by desiccation). The opening was closed with a dry cotton plug. Mortalities were recorded daily until the death of the last individual. For each developmental temperature and species, we ran 10 independent trials in both desiccation and starvation resistance experiments. We performed all the experiment at 23° C and 70% humidity.

### Data analysis

All analyses were conducted in R version 4.3.1 (R Development Core Team, 2023) and the Paleontological Statistics (PAST) version 3.12 (Hammer et al., 2001) software. To visualize the divergence of the CHCs produced by each species across the different temperatures, we used the nonmetric multidimensional scaling (NMDS) with the Bray-Curtis dissimilarity distance (Ricotta and Podani, 2017). To compare the normalized intensity of each species most abundant CHCs across the different developmental temperatures and between sex, we ran the two-way analysis of variance (ANOVA) followed by the Student-Newman-Keul (SNK) posthoc tests, using the R package called “agricolae” (de Mendiburu, 2023). We used the Chi-square test to compare the proportion of mated flies across the rearing temperature and species. To analyze mating duration in relation to rearing temperature and species, we conducted a two-way ANOVA followed by SNK posthoc tests. We used the same test to see whether the fertility, developmental time and body weight (fresh and dry) data varied across the different developmental times and species. We executed Kaplan–Meier survival analysis utilizing functions from the “survival” R package (Therneau, 2023), including *survfit()* and *survdiff()*, in addition to the *pairwise_survdiff()* function available in the “survminer” R package (Kassambra et al., 2021). This analysis was conducted to examine variations in the survival of three drosophilid species concerning different rearing temperatures and species, following exposure to either desiccation or food deprivation. Also, the same analysis was used to compared the longevity data across the different rearing temperature and species. To analyse the median lethal time data from the desiccation and starvation resistance bioassays as well as the longevity assays, we employed a two-way ANOVA, subsequently conducting SNK posthoc tests. Before running the ANOVA test, the normality (Shapiro test: *P>0.05*) and the homoscedasticity (Bartlett test: *P>0.05*) assumptions were verified. Statistical results were considered significant when *P < 0.05*.

## Results

### Different rearing temperatures affect the CHC composition in *D. ezoana*, *D. novamexicana* and *D. virilis*

Rearing temperature significantly affected the composition of cuticular hydrocarbons (CHCs) in the three drosophilid species (Fig. 2). The non-metric multidimensional scaling plots clearly segregated the body extract samples of *D. ezoana* (Fig. 2A), *D. novamexicana* (Fig. 2B), and *D. virilis* (Fig. 2C) based on rearing temperatures. In *D. ezoana*, individuals developed at 15 and 20°C had CHCs with higher peak areas, as opposed to those developed at 25 and 30°C (Fig. 2A). This was the case for heneicos-1-ene (male-specific), 2-heptacosanol, (Z)-13-docosen-1-ol, 2-methyl octacosane, and acetate octadecen-1-ol. In contrast, the change in CHC peaks with increasing rearing temperature was not linear in *D. novamexicana* (Fig. 2B). Compared to individuals developed at 15 and 20°C, those from 25 followed by 30°C produced more (Z)-9-tricosene (Fig. 2B). Furthermore, in a sex-dependent manner, individuals that developed at 15, 20, and 30°C produced more 1-heptacosanol and 2-methyl-octacosane than those from 25°C. While cis-9-eicosen-1-ol was highly produced by *D. novamexicana* developed at 30°C, nonacosanal was highly produced by individuals developed at 15 and 20°C. The production of 2-methyl-triacontane significantly increased with rearing temperature. Also, in *D. virilis*, some CHC peaks did not undergo a linear reduction with increasing rearing temperature (Fig. 2C). For instance, the peaks of 2-methyltetracosane, (Z)-12-pentacosene, 1-heptacosanol and (Z)-13-docosen-1-ol were significantly reduced, while those of 2-methyl-octacosane and nonacosanal were higher in individuals developed at 25°C.

**Figure 1.**
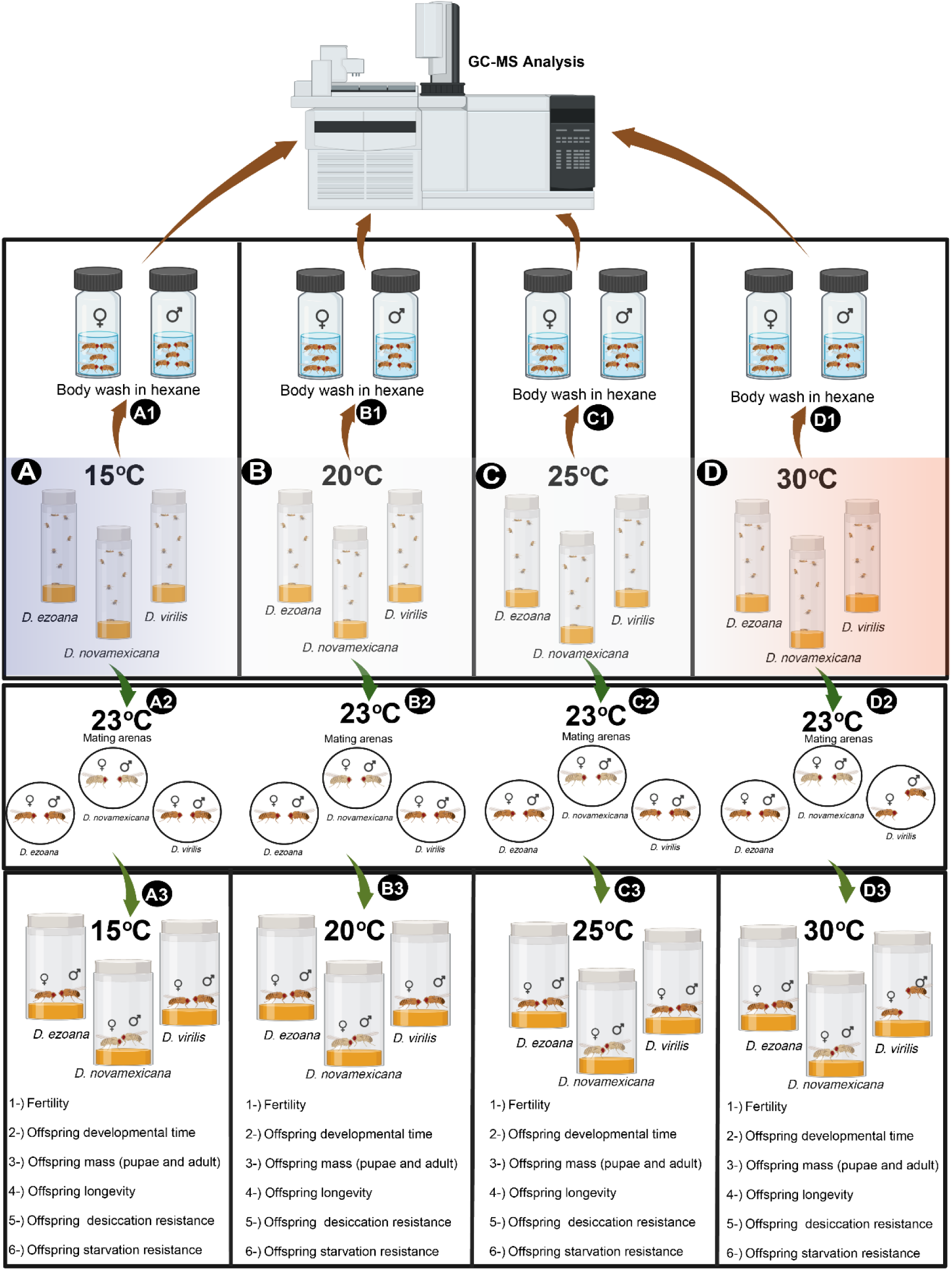
Diagram of the experimental design. (Conceived on <u>biorender.com</u> by Steve B. S. Baleba). *Drosophila ezoana*, *D. novamexicana*, and *D. virilis* adults were reared at temperatures of 15°C (**A**), 20°C (**B**), 25°C (**C**), and 30°C (**D**). The adults were then separated by sexes and soaked in hexane for extraction [(A1), (B1), (C1), and (D1)]. The obtained hexane extracts were subsequently injected into the gas chromatography-mass spectrometry machine. Concurrently, the mating behavior of these adult flies was observed for 1 hour in a mating arena at 23°C [(A2), (B2), (C2), and (D2)]. Mated flies were returned to their respective temperature conditions [(A3), (B3), (C3), and (D3)], and their fertility was assessed. Additionally, developmental time, mass (pupae and adults), longevity, desiccation resistance, and starvation resistance of the offspring produced by these mated flies were recorded.

**Figure 2.**
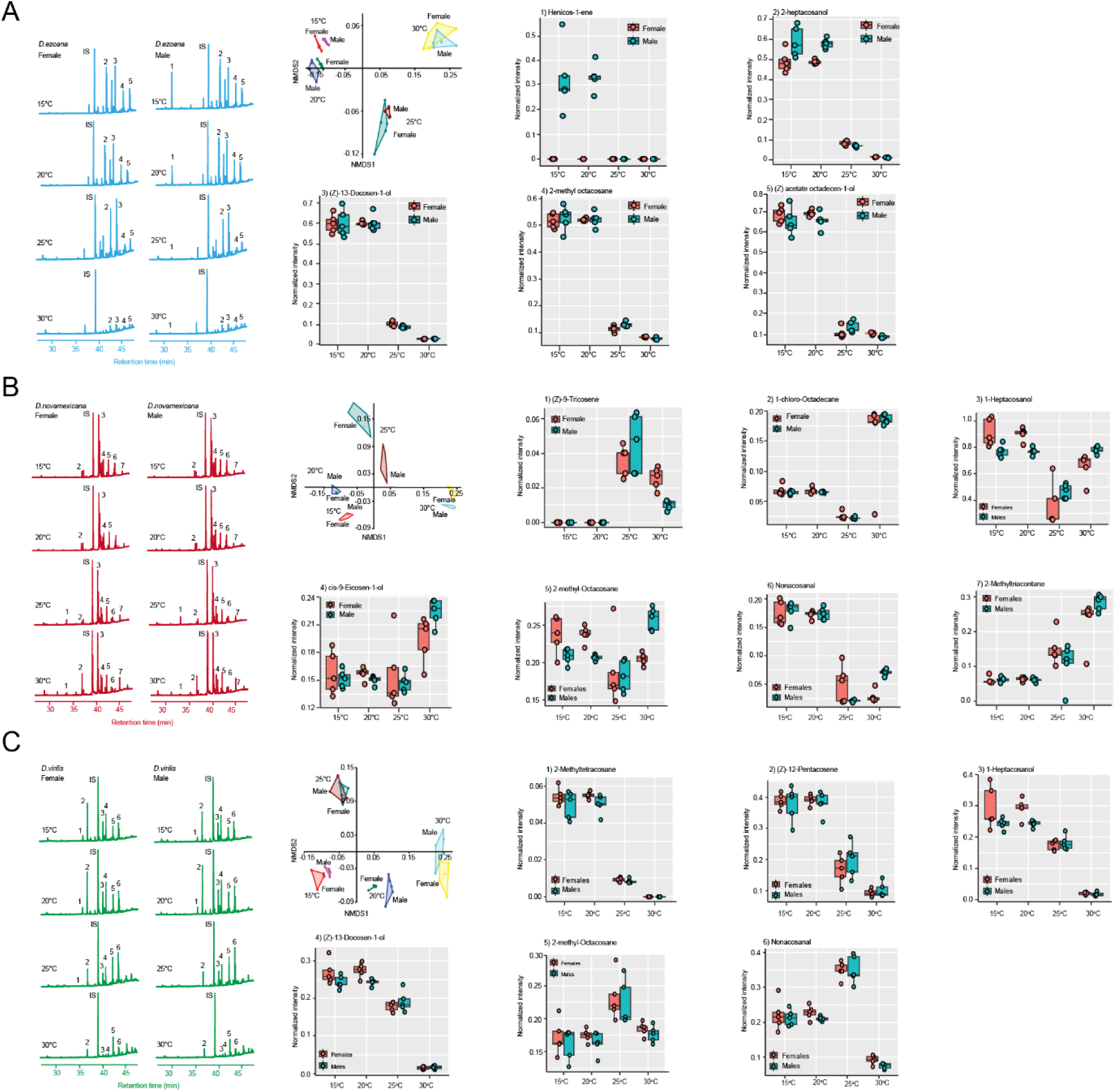
The cuticular hydrocarbon (CHC) composition of the three *Drosophila* species change in function of their rearing temperature. (**A**) *D. ezoana* CHC composition when developed at 15°C, 20°C, 25°C, and 30°C. (**B**) *D. novamexicana* CHC composition when developed at 15°C, 20°C, 25°C, and 30°C. (**C**) *D. virilis* CHC composition when developed at 15°C, 20°C, 25°C, and 30°C. On each figure panel, we present GC-MS chromatogram from different body wash samples, Non-Metric Multidimensional Scaling plot grouping body wash samples based on the rearing temperature, and boxplots illustrating the variation in normalized intensity of the most predominant CHCs across the body wash samples. In each boxplot, the ends of boxplot whiskers represent the minimum and maximum values of all the data and dots show individual data points.

### Higher rearing temperatures leads to reduced mating

Having shown that increased rearing temperatures significantly affect cuticular hydrocarbon compositions in *D. ezoana*, *D. novamexicana* and *D. virilis*, we next aimed to see whether this would affect the mating behavior of these flies, as some of the CHCs are known to mediate sexual communication (Khallaf et al., 2021). We found a significant change in both mating percentage and duration in flies reared at different temperatures (Fig. 3). In *D. ezoana*, individuals developed at 20° C mated frequently, while those developed at 30° C mated significantly less often (Fig. 3A, left). Also, flies developed at 30° C showed a shorter mating duration (Fig. 3A, right). In *D. novamexicana*, individuals developed at 20 and 25° C displayed the highest mating percentage as opposed to those developed at 15 and 30° C (Fig. 3B, left). Here also, individuals developed at 30° C exhibited a shorter mating duration when compared to those developed at 15, 20 and 25° C (Fig. 3B, right). In *D. virilis*, flies developed at 20 and 25° C again showed a higher mating percentage as compared to those developed at 15 and 30° C (Fig. 3C, left). When we compared the mating percentage across the species, we observed that in contrast to *D. novamexicana* and *D. virilis*, *D. ezoana* showed a higher propensity to mate when developed at 15 and 20° C than when they had developed at 25 and 30° C (Supplementary Fig. 1A). We also observed that at both 15 and 30° C, *D. novamexicana* had a shorter mating duration (Supplementary Fig. 1B).

**Figure 3.**
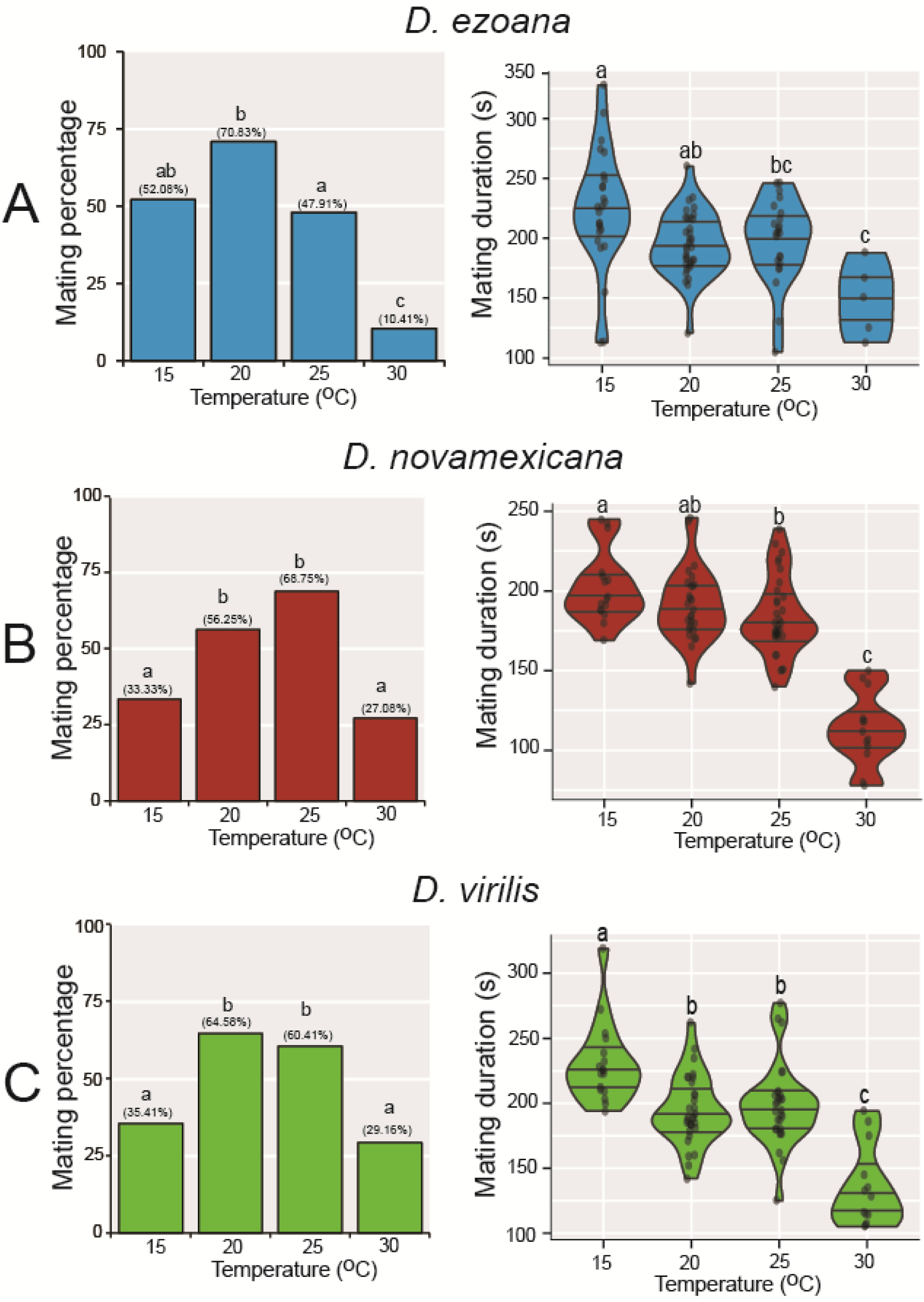
Rearing temperatures affect the mating behavior of the three *Drosophila* species. (**A**) Left, bar graphs showing the variation of mating success in *D. ezoana* when reared at 15°C, 20°C, 25°C, and 30°C. Right, violin plots illustrating the change of the mating time of *D. ezoana* across the different rearing temperature. (**B**) Left, bar graphs depicting the variation of mating success in *D. novamexicana* when developed at 15°C, 20°C, 25°C, and 30°C. Right, violin plots illustrating the change of the mating time of *D. novamexicana* across the different rearing temperature. (**C**) Left, bar graphs depicting the variation of mating success in *D. virilis* when developed at 15°C, 20°C, 25°C, and 30°C. Right, violin plots illustrating the change of the mating time of *D. virilis* across the different rearing temperature. Dots on each violin plot indicate data points from each replicate. Distinct letters on each graph denote significant differences (Mating percentage: Chi-square test; Mating duration: ANOVA followed by the SNK posthoc tests).

### Developmental temperature affects life history traits

After establishing the impact of rearing temperatures on the mating behavior of the three drosophilid species, our next objective was to explore whether the treatments also influenced the fitness of the ensuing offspring. We first assessed the ability of the flies to produce viable adult offspring, a measure of fertility. Across the three species, we observed significant variations in fertility corresponding to changes in rearing temperature (Fig. 4A). Specifically, in *D. ezoana* (Fig. 4A, left), *D. novamexicana* (Fig. 4A, middle), and *D. virilis* (Fig. 4A, right), females developed and maintained at 15°C and 30°C produced fewer adult offspring than those developed and maintained at 20°C and 25°C. Additionally, it was apparent that at each rearing temperature, *D. novamexicana* exhibited lower fertility compared to *D. ezoana* and *D. virilis* (Fig. 4B).

**Figure 4.**
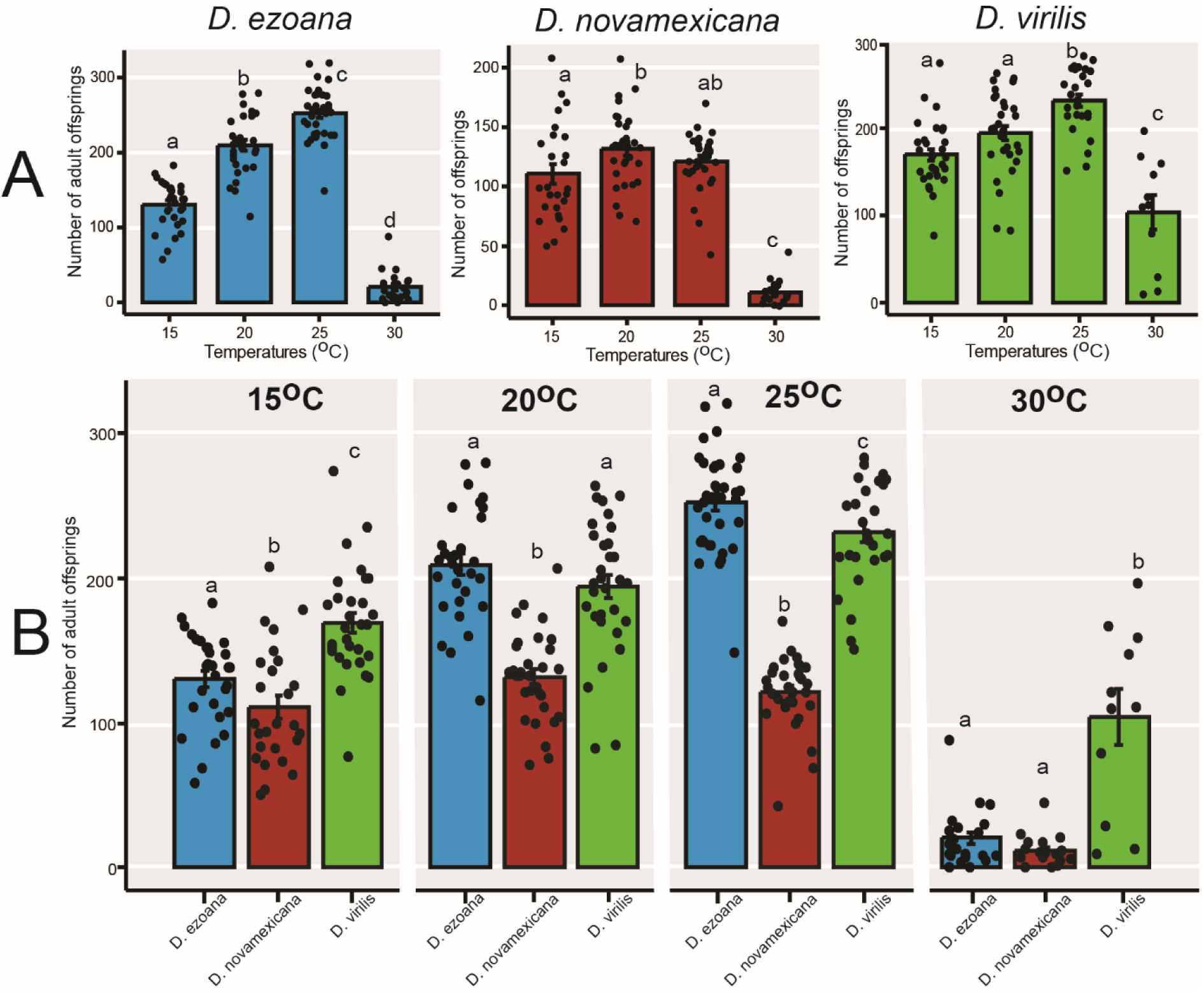
Rearing temperatures influences the fertility of the three *Drosophila* species. (**A**) Bar graphs showing the mean number of offspring produced by females of *D. ezoana* (left), *D. novamexicana* (middle) and *D. virilis* (right) when developed and maintained at 15°C, 20°C, 25°C, and 30°C. (**B**) Bar graphs displaying the mean number of offspring produced by females of *D. ezoana*, *D. novamexicana* and *D. virilis* for each rearing temperature. Dots on each bar graph show individual data point. Error bars indicate standard error of the mean (SEM). Bars with different letters are significantly different from each other (ANOVA followed by the SNK posthoc tests; *P* < 0.05, n = 20-40).

However, prior to these offspring reaching adulthood, we observed significant changes in their developmental time (Fig. 5A) and pupal weight (Fig. 5B) in response to rearing temperature. In *D. ezoana* (Fig. 5A, left), *D. novamexicana* (Fig. 5A, middle), and *D. virilis* (Fig. 5A, right), offspring developed at 15°C required a longer time to reach adulthood compared to those developed at 20, 25 and 30°C. When considering the species as a whole, it was apparent that at 15°C, *D. ezoana* offspring, followed by those of *D. virilis*, exhibited a shorter developmental time compared to those of *D. novamexicana* (Supplementary Fig. 2A). Moreover, at 30°C, *D. virilis* offspring reached adulthood at a faster rate than their counterparts from *D. ezoana* and *D. novamexicana*.

**Figure 5.**
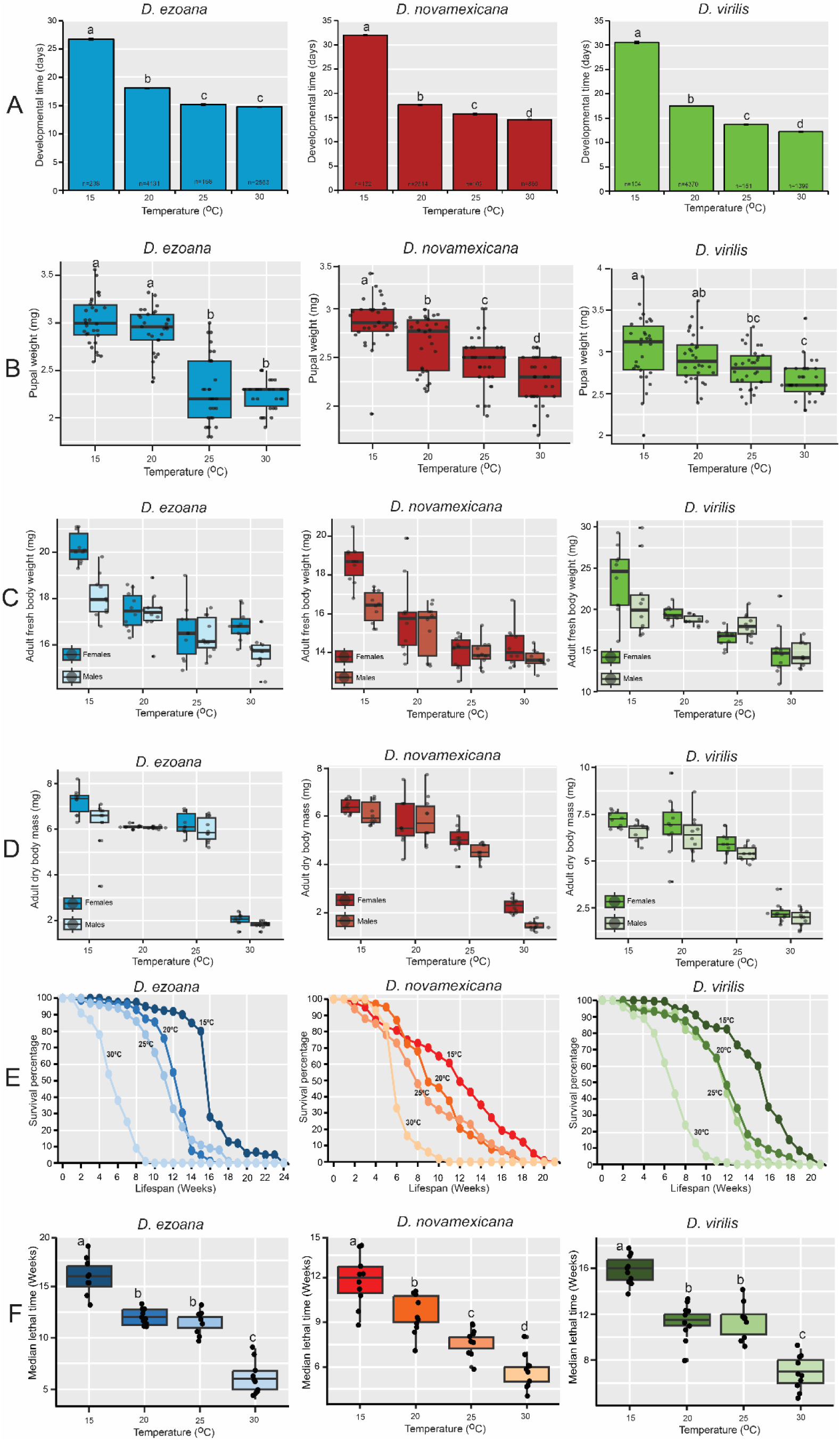
Rearing temperatures affect the life history parameters of the offspring produced by the three *Drosophila* species. (**A**) Bar graphs showing the mean developmental time of the offspring of *D. ezoana* (left), *D. novamexicana* (middle) and *D. virilis* (right) when developed at 15°C, 20°C, 25°C, and 30°C. Error bars indicate standard error of the mean (SEM). Bars with different letters are significantly different from each other (ANOVA followed by the SNK posthoc tests; *P* < 0.05). (**B**) Boxplots depicting the pupae weight of *D. ezoana* (left), *D. novamexicana* (middle) and *D. virilis* (right) when developed at 15°C, 20°C, 25°C, and 30°C. (**C**) Boxplots depicting the fresh adult (female and male) body weight of *D. ezoana* (left), *D. novamexicana* (middle) and *D. virilis* (right) when developed at 15°C, 20°C, 25°C, and 30°C. (**D**) Boxplots depicting the dry adult (female and male) body weight of *D. ezoana* (left), *D. novamexicana* (middle) and *D. virilis* (right) when developed at 15°C, 20°C, 25°C, and 30°C. (**E**) Kaplan-Meier curves showing the variation of the longevity of adult offspring produced by *D. ezoana* (left), *D. novamexicana* (middle) and *D. virilis* (right) when developed at 15°C, 20°C, 25°C, and 30°C. (**F**) Boxplots illustrating the change of the median lethal time of adult offspring produced by *D. ezoana* (left), *D. novamexicana* (middle) and *D. virilis* (right) when developed at 15°C, 20°C, 25°C, and 30°C. In each boxplot, the ends of boxplot whiskers represent the minimum and maximum values of all the data and dots show individual data points. Box plots with different letters are significantly different from each other (ANOVA followed by the SNK posthoc tests; *P* < 0.05, n = 10).

Furthermore, we observed that pupae of *D. ezoana* (Fig. 5B, left), *D. novamexicana* (Fig. 5B, middle), and *D. virilis* (Fig. 5B, right) displayed increased weight when developed at 15°C and 20°C, as opposed to those developed at 25 and 30°C. When comparing across species, it became evident that pupae from *D. ezoana* and *D. virilis* exhibited greater weight at 15 and 20°C, while at 25°C and 30°C, only *D. virilis* pupae demonstrated a higher weight (Supplementary Fig. 2B).

When adults emerged from these pupae, we observed a significant decrease in their fresh (Fig. 5C) and dry (Fig. 5D) body weights as developmental temperatures increased. The fresh body mass of adult offspring from *D. ezoana* (Fig. 5C, left), *D. novamexicana* (Fig. 5C, middle), and *D. virilis* (Fig. 5C, right) was highest in individuals developed at 15°C (with females outweighing males), followed by those at 20, 25, and 30°C. Notably, at all developmental temperatures except 30°C, adult offspring of *D. novamexicana* were lighter in weight compared to that of *D. ezoana* and *D. virilis* (Supplementary Fig. 2C). Additionally, adult offspring of *D. ezoana* (Fig. 5D, left), *D. novamexicana* (Fig. 5D, middle), and *D. virilis* (Fig. 5D, right) developed at 15°C and 20°C had heavier dry body weights than those developed at 25°C and 30°C. When comparing across species, adult offspring of *D. novamexicana* developed at 15, 20 and 25°C, consistently exhibited lower dry body weights than those from *D. ezoana* and *D. virilis* (Supplementary Fig. 2D).

We also observed that the longevity of adult offspring from *D. ezoana* (Fig. 5E, left), *D. novamexicana* (Fig. 5E, middle), and *D. virilis* (Fig. 5E, right) decreased significantly with increase of developmental temperatures. In the three species, adult offspring developed at 15, 20 and 25°C lived longer than those developed at 30°C. Specifically, in the case of *D. ezoana* (Fig. 5F, left), *D. novamexicana* (Fig. 5F, middle), and *D. virilis* (Fig. 5F, right), half of the offspring developed at 15°C survived for a significantly longer duration compared to those raised at 30°C. Furthermore, when comparing across species, we observed that adult offspring of *D. novamexicana* in general had a shorter longevity than those of *D. ezoana* and *D. virilis* (Supplementary Fig. E and F).

### Changes in rearing temperature affect resistance to desiccation and starvation

As a rise of temperature leads to increased water loss and, subsequently, reduction in food availability, we assessed the desiccation and starvation resistance in the adult offspring of the three drosophilid species as developed at 15, 20, 25 and 30°C. The results from the desiccation resistance experiment highlighted the significant impact of rearing temperature on the ability of the adult offspring from these species to endure and survive prolonged dry conditions (Fig. 6). In the case of *D. ezoana*, adult offspring developed at 20°C and 25°C exhibited markedly higher resistance to silica gel exposure compared to those raised at 15°C and 30°C (Fig. 6A, left), with a delayed mortality observed in half of the population (Fig. 6A, right). In *D. novamexicana*, adult offspring developed at 25°C and 30°C demonstrated greater desiccation resistance compared to those raised at 15°C and 20°C (Fig. 6B, left and right). Similarly, adult offspring of *D. virilis* developed at 25 and 30°C exhibited greater resistance to silica gel exposure than those developed at 15°C and 20°C (Fig. 6C, left and right). When comparing desiccation resistance across species, it became apparent that at developmental temperatures of 15°C (Fig. 6D, left and right), 20°C (Fig. 6E, left and right), 25°C (Fig. 6F, left and right), and 30°C (Fig. 6G, left and right), *D. virilis*, followed by *D. novamexicana*, displayed the highest levels of resistance to desiccation.

**Figure 6.**
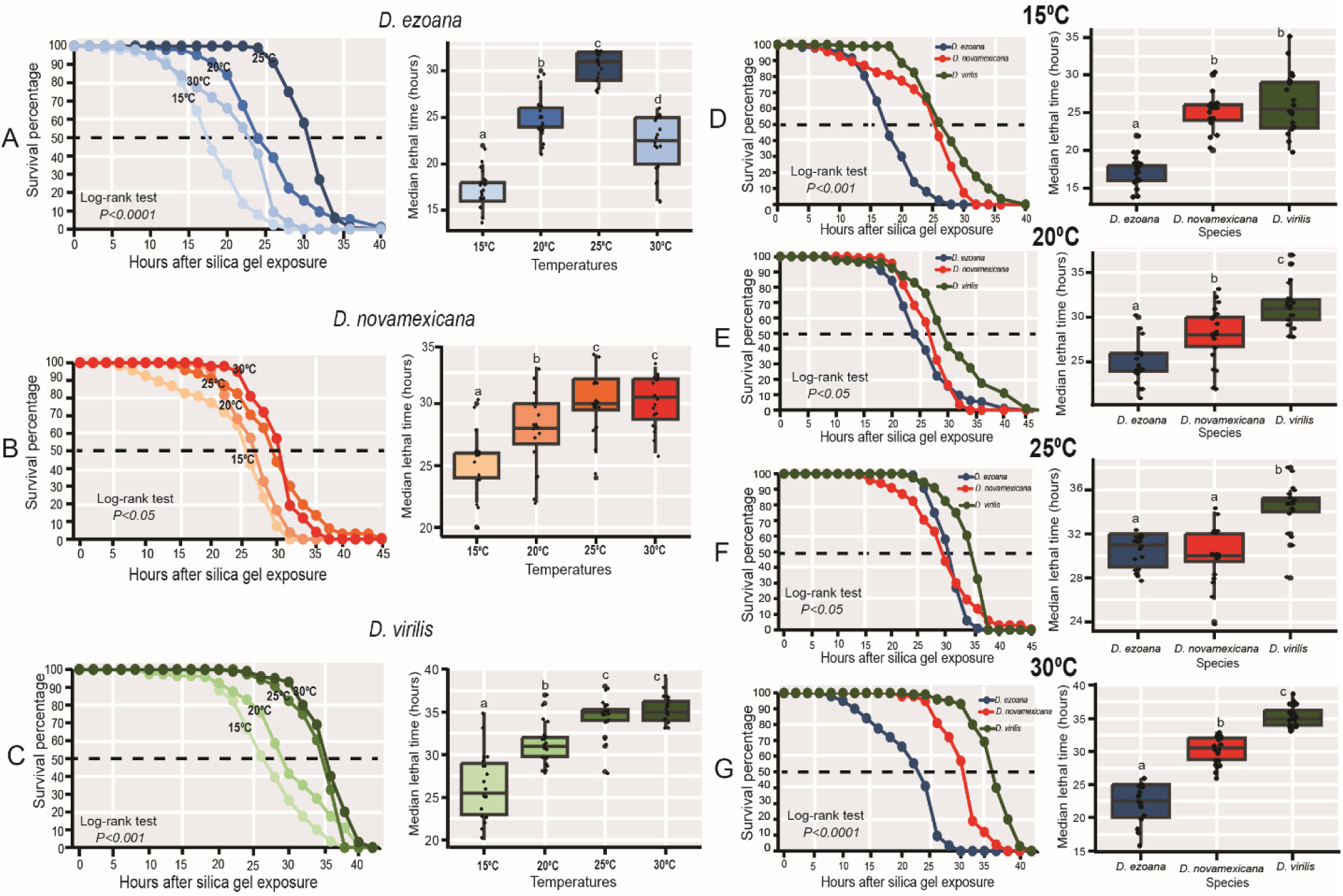
Rearing temperatures influence the desiccation resistance of the offspring produced by the three *Drosophila* species. (**A-B**) Kaplan–Meier curves (left) and Boxplots (right) showing respectively the change of the survivorship (after silica gel exposure) and median lethal time of the offspring of *D. ezoana* (**A**), *D. novamexicana* (**B**) and *D. virilis* (**C**) across the different rearing temperatures (15°C, 20°C, 25°C, and 30°C). (**D-G**) Kaplan–Meier curves (left) and Boxplots (right) illustrating respectively the variation of the survivorship (after silica gel exposure) and the median lethal time across the three *Drosophila* species when developed at 15°C (**D**), 20°C (**E**), 25°C (**F**), and 30°C (**G**). Each boxplot displays the median and whiskers indicate ± 1.5 interquartile range limits. Dots on each box plot show individual data point. Box plots with different letters are significantly different from each other (ANOVA followed by the SNK posthoc tests; *P* < 0.05, n = 10).

The response to food deprivation varied among adult offspring of *D. ezoana*, *D. novamexicana*, and *D. virilis* when reared at different temperatures (Fig. 7). In the case of *D. ezoana*, individuals that developed at 15°C demonstrated greater resistance to food deprivation compared to those developed at 20, 25 and 30°C, with the latter group showing the highest susceptibility to starvation (Fig. 7A, left, right). For *D. novamexicana*, adult offspring developed at 20 and 25° C were more resistant to starvation than those from 15 and 30° C (Fig. 7B, left, right). In *D. virilis*, adult offspring developed at 15 and 20° C were more starvation resistant compared to those developed at 25 and 30° C (Fig. 7C, left, right). When compared across species, only adult offspring developed at 15°C showed similar starvation resistance across the three species (Fig. 7D, left, right). At developmental temperature of 20 (Fig. 7E, left, right), 25 (Fig. 7F, left, right), and 30° C (Fig. 7G, left, right), adult offspring produced by *D. novamexicana* followed by those of *D. virilis* exhibited greater starvation resistance as compared to their counterpart produced by *D. ezoana*.

**Figure 7.**
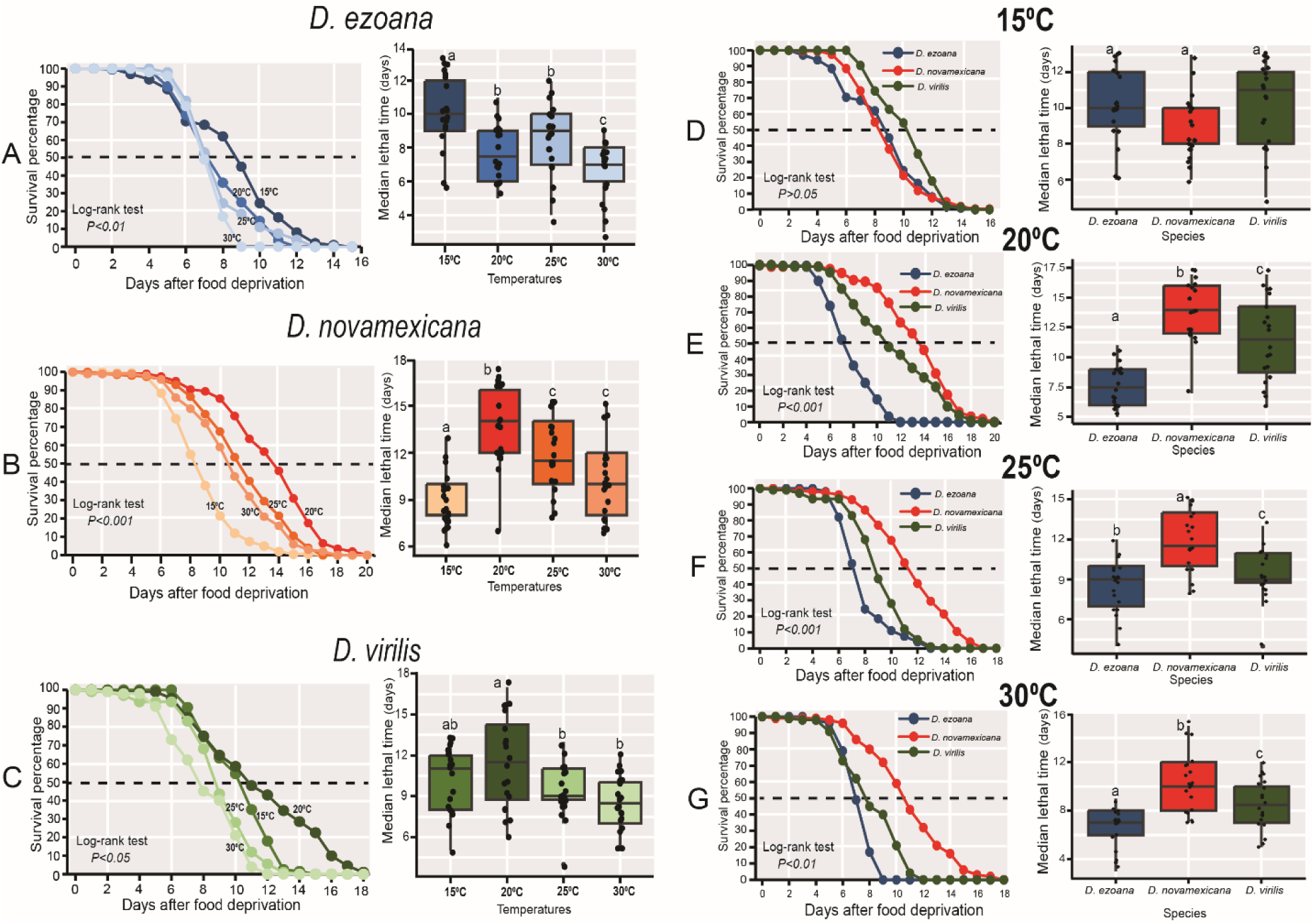
Rearing temperature influence the starvation resistance of the offspring produced by the three *Drosophila* species. (**A-B**) Kaplan–Meier curves (left) and Boxplots (right) showing respectively the change of the survivorship (after food deprivation) and median lethal time of the offspring of *D. ezoana* (**A**), *D. novamexicana* (**B**) and *D. virilis* (**C**) across the different developmental temperatures (15°C, 20°C, 25°C, and 30°C). (**D-G**) Kaplan–Meier curves (left) and Boxplots (right) illustrating respectively the variation of the survivorship (after food deprivation) and the median lethal time across the three *Drosophila* species when developed at 15°C (**D**), 20°C (**E**), 25°C (**F**), and 30°C (**G**). In each boxplot, the ends of boxplot whiskers represent the minimum and maximum values of all the data and dots show individual data points. Box plots with different letters are significantly different from each other (ANOVA followed by the SNK posthoc tests; *P* < 0.05, n = 10).

## Discussion

Variations in rearing temperatures influence the composition of cuticular hydrocarbons (CHCs) in three *Drosophila* species: *D. ezoana*, found in Arctic regions; *D. novamexicana*, native to warm, desert environments and *D. virilis*, with a cosmopolitan distribution. Temperature changes in parallel affect mating behaviour and fertility, and other life history traits in all three species, and also influence desiccation and starvation resistance of the offspring.

We observed distinct variations in cuticular hydrocarbon (CHCs) profiles among adult flies of the three drosophilids species when developed at varying temperatures. In the cold-adapted *D. ezoana*, we observed a consistent decline in the quantity of all CHCs as the developmental temperature increased. Such a trend was not observed in *D. novamexicana* and *D. virilis* (Fig. 2). For instance, *D. novamexicana* individuals developed at 15 and 20°C produced lower amounts of (Z)-9-tricosene and 2-methyl triacontane compared to those developed at 25 and 30°C. However, at the same temperature (15 and 20°C), they synthetized higher amounts of 1-heptacosanol and nonacosanal. Similarly, in *D. virilis*, adults developed at 15 and 20°C produced higher quantities of 2-methyl tetracosane, (Z)-pentacosene, (1)-heptacosanol, and (Z)-13-docosen-ol, while displaying lower levels of 2-methyl-octacosane and nonacosanal. These observations highlight that temperature-dependent shifts in CHC profiles are species-specific. In fact, numerous studies conducted at both the intra– and interspecific levels in insects have consistently confirmed the presence of this plasticity in CHC production in response to environmental stress (…..). Furthermore, comparative studies across various drosophilid species have revealed that species from warmer climates tend to produce a higher proportion of methyl-branched CHCs, in contrast to those from cooler regions. In our case, we observed that, in addition to the common 2-methyl-octadecane produced by all three drosophilid species, the desert species, *D. novamexicana* and the cosmopolitan species *D. virilis* produced 2-methyl-triacontane and 2-methyl-tetracosane, respectively. Wang et al. (2022) found a similar trend in *Drosophila mojavensis*, a desert species known for producing more methyl-branched CHCs. This phenomenon is not unique to *Drosophila* flies, as other desert insect species such as the stink beetle *Eleodes armata* (Hadley, 1977), the locust *Schistocerca gregaria* (Heifetz et al., 1998), the ants *Cataglyphis niger* (Soroker and Hefetz, 2000), and *Pogonomyrmex barbatus* (Johnson and Gibbs, 2004) also exhibit high proportions of methyl-branched CHCs. Wang et al. (2023) discovered that the ability of *D. mojavensis* to synthesize these long methyl-branched cuticular hydrocarbons is linked to genetic variations in a fatty acyl-CoA elongase gene (*mElo*). However, it is important to note that their findings were based on *D. mojavensis* developed at room temperature. Conducting additional studies using flies developed at different temperatures (as we did) to assess the expression levels of the *mElo* gene will provide a deeper understanding of how the temperatures experienced during preimaginal development influence CHCs production in flies.

As well as the significant changes in the compositions of CHCs, we observed alterations in the mating behavior of the three drosophilid species in response to temperature variations during their development. In all three species, individuals developed at 15°C and 30°C exhibited lower mating success compared to those developed at 20°C and 25°C. This difference can be attributed to the impact of temperature on behavior, morphology, and physiology (García-Roa et al., 2020). Insects that develop at extreme temperatures often experience adverse alterations in their biology, rendering them less competent in successful copulation. Lower temperatures tend to slow down metabolic processes, leading to reduced activity levels and increased time spent seeking warmth or shelter, which in turn limits their opportunities to mate (Sinclair et al., 2003). Conversely, insects developing at higher temperatures exhibit higher metabolic rates, which can make them more agitated and stressed, potentially disrupting their mating behaviors (Hannigan et al., 2023). Extreme temperatures can also influence the production, release, and detection of pheromones, affecting mate location and successful copulation (Boullis et al., 2016). For example, henicos-1– ene (male specific pheromone) and (Z)-12-pentacosene from *D. ezoana* and *D. virilis* respectively are produced in lower amounts by individuals developed at 30°C that exhibit lower mating success. Conversely, insects developed at more optimal temperatures tend to produce pheromones in quantities sufficient for successful mating. For instance, in *D. novamexicana*, individuals developed at 25°C that show higher mating percentage also produce significantly higher amounts of (Z)-9-tricosene.

When comparing across species, we observed that the mating success of *D. ezoana* as compared to that of *D. novamexicana* and *D. virilis* was higher when they developed at 15 and 20°C. However, developing at 25 and 30°C, this species showed lower mating percentage. This implies that the cold-adapted species (*D. ezoana*) may have evolved temperature-dependent reproductive traits that are more effective in colder conditions. Conversely, these traits are more evolved under warmer conditions in the warm-adapted (*D. novamexicana*) and the cosmopolitan (*D. virilis*) species. In *D. melanogaster* heat resistant flies mate more at elevated temperature than the heat-susceptible ones (Stazione et al. 2023). Using 12 species of *Drosophila* species, Schnebel and Grossfield (1984) observed that changes in temperature-dependent mating reflect differences in their thermal adaptation and geographic distributions.

In addition to the observed changes in the mating success of the three drosophilid species, we also noted a substantial reduction in mating duration with the rise in developmental temperature. This change is likely a result of various factors, including changes in developmental speed, metabolic rates, behavior and environmental conditions. Collectively, these factors contribute to shorter mating durations in insects exposed to higher temperatures during their development. The three drosophilid species had shorter developmental times (Fig. 5A) and lifespan (Fig. 5E, F) when developed at higher temperatures. As elucidated by Liang et al. (2019), under such a scenario, insects may have less time available for mating before they reach the end of their natural lifespan. Furthermore, the increased metabolic activities in insects when developed at higher temperatures can result in quicker exhaustion of energy reserves, reducing the duration of mating activities (Pilakouta and Baillet, 2022). Moreover, insects developed at higher temperature are perpetually exposed to heat and desiccation stress. To cope with this, Pérez-Staples et al. (2018) explain that insects tend to allocate fewer resources to reproduction and invest more in behavioral and physiological strategies aimed at bolstering their resilience to environmental stressors. This negative relationship between temperature and mating duration has also been observed in studies involving different insect species, such as the Japanese beetle *Popillia japonica* (Saeki et al., 2005), bugs including *Anthocoris tomentocus*, *A. whitei*, *A. nemoralis* (Horton et al. 2002), and the beetle *Callosobruchus chinensis* (Katsuki and Miyatake, 2009).

We observed significant changes in the fertility among the three drosophilid species in response to variations in their developmental temperatures. Specifically, individuals reared at 30°C and 15°C exhibited reduced production of adult offspring compared to those developed at 20°C and 25°C. These findings align with Cohet and David (1978), who demonstrated that *D. melanogaster* females raised at various temperatures (ranging from 12°C to 32°C) produced more adult offspring when reared at 21°C and 25°C. Similar outcomes were also evident in *D. suzikii* (Schlesener et al., 2020). Our study further revealed that flies reared at 15°C and 30°C displayed a decreased mating percentage, providing a potential explanation for the reduced adult offspring at these extreme temperatures. Additionally, extreme temperature conditions can adversely affect sperm production and viability in male insects, leading to diminished fertility due to lower fertilization success. For example, Gandara and Drummond-Barbosa (2023) observed that *D. melanogaster* males reared at 29°C produced fewer and lower-quality sperm, resulting in less efficient egg fertilization. Ovariole number emerged as another critical factor directly impacting fertility. Klepsatel et al. (2019) found that *D. melanogaster* females developed at 17°C and 29°C exhibited lower ovariole numbers and displayed decreased fertility compared to those reared at 25°C. Moreover, insect eggs are highly sensitive to both high and low temperatures, which can disrupt their development, reduce hatchability and result in the production of non-viable offspring (Cavieres et al., 2018). Among the three species, *D. novamexicana* stood by producing fewer adult offspring. Notably, *D. novamexicana* females displayed a smaller fresh body weight in comparison to *D. ezoana* and *D. virilis* females (see Supplementary Fig. 2C).). According to Berger et al. (2008) female body size is a reliable predictor of fertility in general. Larger female insects have the capacity to accumulate more resources and convert them into eggs, often yielding more viable adult offspring. This positive correlation between body size and fertility is also found in various insect species including the moth *Streblote panda* (Calvo and Molina, 2005), the beetl *Callosobruchus chinensis* (Yanagi and Tuda, 2012; Yanagi and Miyatake, 2002), and the flies *Drosophila malerkotliana* and *D. bipectinate* (Krishna and Hegde, 1997).

We observed that as the developmental temperature increased, the females of the three drosophilid species produced adult offspring characterized by shorter development periods, decreased pupal and adult weights (both fresh and dry) and reduced longevity. Indeed, insect development is governed by enzymes whose activities increase in warmer conditions, consequently accelerating crucial chemical reactions essential for growth, molting, and overall development (Arcus et al., 2016; Shinoda and Itoyama, 2003). The observed reduction in size in our study can be attributed to accelerated development at higher temperatures, which offer the flies less time for feeding, growth, and nutrient storage before their pupation and emergence (Hoover and Marks, 2021). Like in almost all ectotherms, our three drosophilid species follow the temperature-size rule (TSR) stating that in insects, an increase in developmental temperature leads to a decrease in final adult size (Atkinson, 1994; Ghosh et al., 2013). Using *D. melanogaster*, Li and Gong (2015) demonstrated that development under cold conditions increases the production of insulin-like peptides leading to an increase of body size. Furthermore, apart from the reduction in body size, our study also uncovered a decrease in the lifespan of adult offspring produced by these three drosophilid species. A study using *D. melanogaster* from different genetic background showed that as temperature increases, the metabolic rate of insects rises, leading to higher energy expenditure and faster depletion of stored resources, thereby shortening their lifespan (Mołoń et al., 2020). Additionally, elevated temperatures can raise the production of detrimental reactive oxygen species (ROS) within the insect’s body, potentially causing damage to cellular components and leading to oxidative stress, which is linked to both aging and diminished longevity (Jena et al., 2013; Paital, 2016).

When compared at the species level, *D. novamexicana* exhibited smaller body size and decreased longevity in comparison to *D. ezoana* and *D. virilis* (Supplementary Fig. 2). In many insects including moths (Holm et al., 2016), cockroaches (Badwan and Harper, 2021), and mosquitoes (Barreaux, 2018; Christiansen-Jucht et al., 2014; Takken et al., 1998), smaller individuals tend to have shorter lifespans. This phenomenon can be attributed to the fact that smaller insects tend to be less efficient at accumulating food reserves and typically exhibit higher metabolic rates. Consequently, these factors can lead to accelerated aging and a subsequent reduction in their overall lifespan (Promislow et al., 2022).

We found an intra and interspecific variation of desiccation resistance in the three drosophilid species when developed at different temperatures. The flies exhibited lower resistance to water loss when developed at 15°C. Adult insects emerging from development at lower temperatures, consistently exhibit increased susceptibility to desiccation stress due to reduced prior exposure to dehydration stress (Bong et al., 2021a; Yoder et al., 2011). On the other hand, a non-lethal exposure to desiccation stress during development increases resistance to subsequent exposures (Kellermann et al., 2018). This might explain why, when developed at 25 and 30°C, the flies showed high desiccation resistance. When compared across species, the cold-adapted species (*D. ezoana*), was less resistant to water loss compared to the warm-adapted (*D. novamexicana*) and cosmopolitan (*D. virilis*) species. In fact, xeric-adapted and cosmopolitan insects like *D. novamexicana* and *D. virilis* respectively are generally more resistant to desiccation compared with mesic-adapted insects like *D. ezoana* (Bong et al., 2021b; Matzkin et al., 2009a). This is mainly because they produce higher proportion of CHCs as we previously elucidated. Indeed there is a causal association between CHCs and desiccation survival (Foley and Telonis-Scott, 2011), where an increase in desiccation resistance is associated with a high CHCs proportion (Ferveur et al., 2018).

We also observed changes in starvation resistance in response to variation in developmental temperature. In *D. melanogaster*, exposure to low temperatures during preimaginal development diminishes resistance to starvation, whereas higher temperatures enhance it (Jang and Lee, 2018). However, this was not the case in our three species. In particular, the cold-adapted *D. ezoana* and, to some extent, the cosmopolitan *D. virilis* exhibited heightened starvation resistance when developed at 15°C (Fig. 7A and C). Conversely, individuals from the warm-adapted species, *D. novamexicana* (Fig. 7B), displayed reduced resistance to starvation when subjected to the same developmental temperature. When developed at 30°C, *D. ezoana* experienced a significant reduction in starvation resistance compared to *D. novamexicana* and *D. virilis*. Matzkin et al. (2009) observed that cactophilic drosophilid species, adapted to arid or desert regions, tend to withstand longer periods of food deprivation than their fruit-breeding counterparts. Insects in arid or desert regions often experience prolonged periods of food scarcity. They have evolved physiological and behavioral adaptations to reduce metabolic rates, optimize water use, and store energy more efficiently to survive extended periods of starvation (Zhang et al., 2019). This provides a plausible explanation why *D. novamexicana* exhibited the highest starvation resistance. Additionally, earlier research by Da Lage et al. (1990) unveiled that Afrotropical *D. melanogaster* flies, which inhabit hot environments, are approximately twice as resistant to starvation as their temperate counterparts, providing further insight into the complex relationship between developmental conditions and starvation resistance in drosophilid species.

## Conclusion

We studied the impact of rearing temperature on three *Drosophila* species—*D. ezoana*, *D. novamexicana*, and *D. virilis*—across various facets, including cuticular hydrocarbon profiles, mating behavior, fertility, life history traits, and desiccation and starvation resistance. Notably, each species exhibited unique variations in cuticular hydrocarbon profiles in response to different developmental temperatures, a phenomenon consistent with the notion that species from warmer climates tend to produce more methyl-branched CHCs. Mating behavior was significantly affected by developmental temperature, with 15°C and 30°C leading to lower mating success, potentially attributed to alterations in behavior, physiology and pheromone production. Furthermore, flies developed at 15°C and 30°C displayed reduced fertility, influenced by factors such as mating success and duration. The study also revealed that higher temperatures resulted in smaller body sizes and decreased lifespans in adult offspring, driven by accelerated development. In terms of resistance to desiccation and starvation, the warm-adapted and cosmopolitan species demonstrated greater resistance due to their higher proportion of cuticular hydrocarbons in comparison to the cold-adapted species. Taken together, our study underscore the intricate relationship between developmental temperature, ecological adaptation and various life history traits, contributing valuable insights into how changes in environmental factors shape the biology and ecology of distinct species.

## Author contributions

**Steve B. S. Baleba:** Conceptualization, methodology, data curation; formal analysis; visualization; writing – original draft

**Nan-Ji Jiang**: Methodology; visualization; review and editing

**Bill S. Hansson:** Fund acquisition, supervision, methodology, review and editing

## Acknowledgements

We express our gratitude to Angela Lehmann for her invaluable assistance in configuring and setting up the incubators.

## Data availability statement

The data supporting the findings of this study will be accessible on https://figshare.com/ website upon manuscript acceptance

## Supplementary Figures

**Supplementary figure 1.**
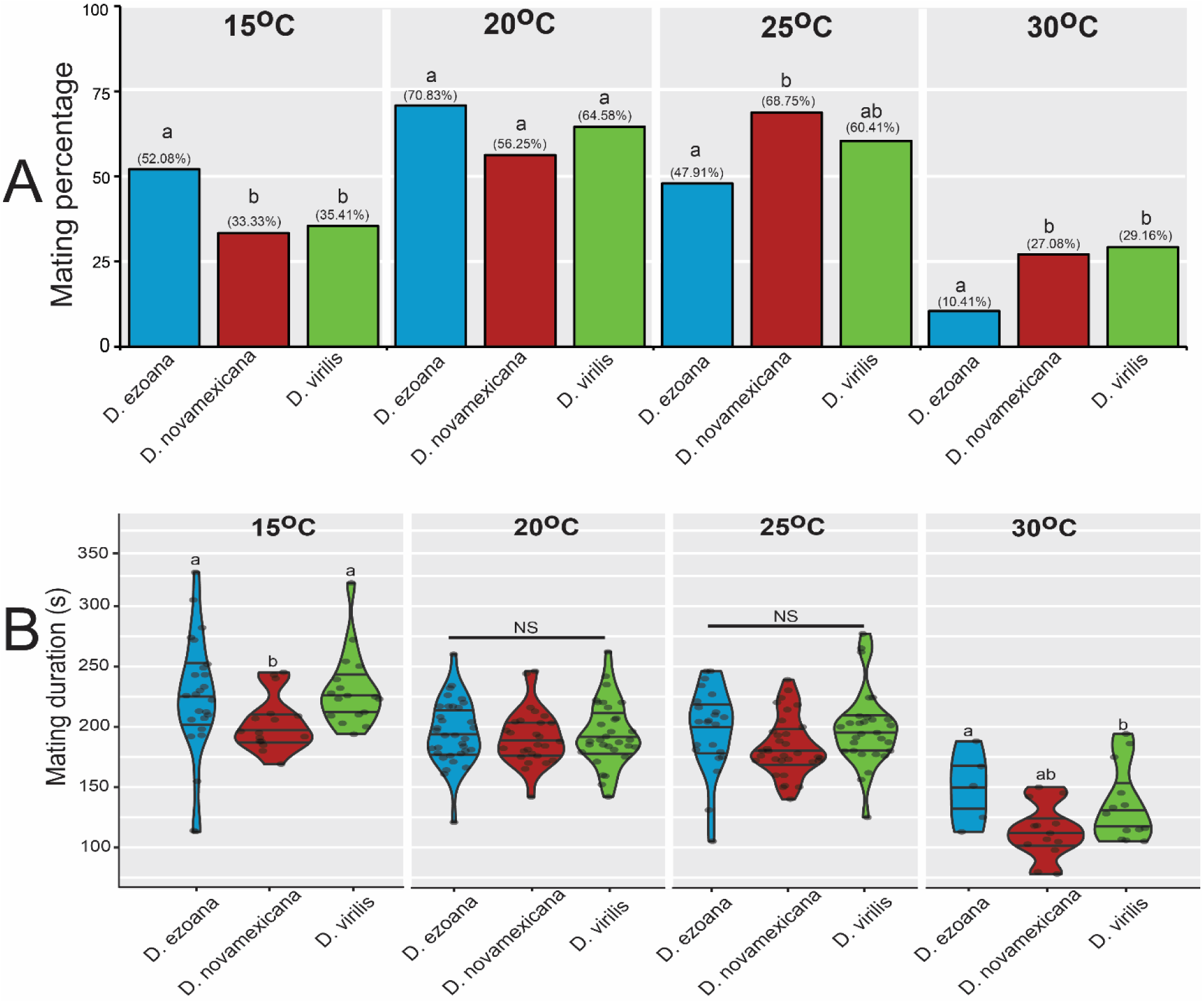
The three *Drosophila* species exhibit distinct mating behavior. (**A**) Bar graphs comparing the mating percentage of *D. ezoana*, *D. novamexicana* and *D. virilis* when developed at 15, 20, 25, and 30°C. (B) Violin plots comparing the mating time of *D. ezoana*, *D. novamexicana* and *D. virilis* when developed at 15, 20, 25, and 30°C. Dots on each violin plot indicate data points from each replicate. Distinct letters on each graph denote significant differences (Mating percentage: Chi-square test; Mating duration: ANOVA followed by the SNK posthoc tests).

**Supplementary figure 2.**
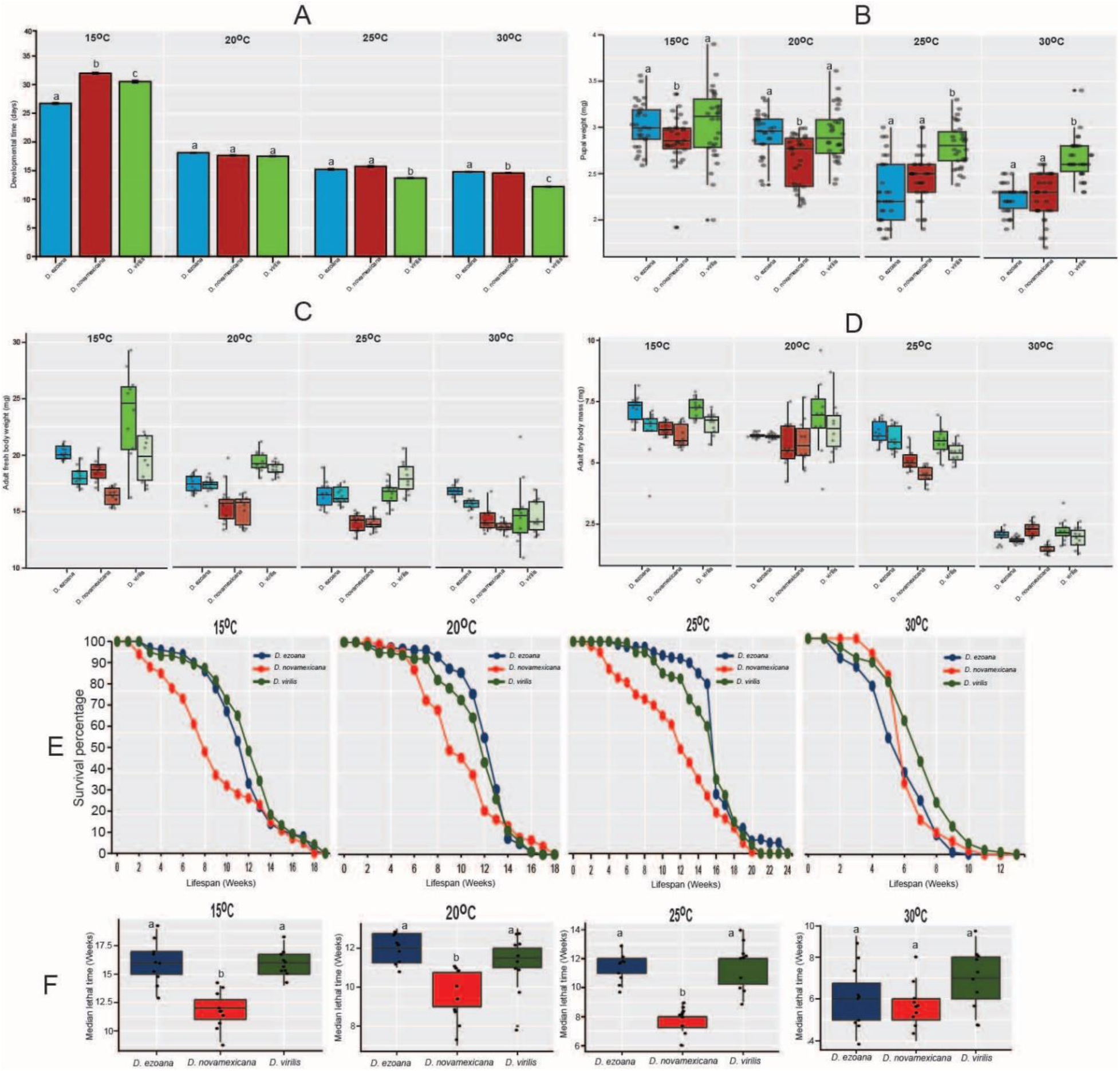
The three *Drosophila* species exhibit different life history traits. (**A**) Bar graphs comparing the mean developmental time of the offspring of *D. ezoana*, *D. novamexicana* and *D. virilis* when developed at 15°C, 20°C, 25°C, and 30°C. Error bars indicate standard error of the mean (SEM). Bars with different letters are significantly different from each other (ANOVA followed by the SNK posthoc tests; *P* < 0.05). (**B**) Boxplots comparing the pupae weight of *D. ezoana*, *D. novamexicana* and *D. virilis* when developed at 15°C, 20°C, 25°C, and 30°C. (**C**) Boxplots comparing the fresh adult (female and male) body weight of *D. ezoana*, *D. novamexicana* and *D. virilis* when developed at 15°C, 20°C, 25°C, and 30°C. (**D**) Boxplots comparing the dry adult (female and male) body weight of *D. ezoana*, *D. novamexicana* and *D. virilis* when developed at 15°C, 20°C, 25°C, and 30°C. (**E**) Kaplan-Meier curves comparing the variation of the longevity of adult offspring produced by *D. ezoana*, *D. novamexicana* (middle) and *D. virilis* when developed at 15°C, 20°C, 25°C, and 30°C. (**F**) Boxplots comparing the change of the median lethal time of adult offspring produced by *D. ezoana*, *D. novamexicana* and *D. virilis* when developed at 15°C, 20°C, 25°C, and 30°C. In each boxplot, the ends of boxplot whiskers represent the minimum and maximum values of all the data and dots show individual data points. Box plots with different letters are significantly different from each other (ANOVA followed by the SNK posthoc tests; *P* < 0.05, n = 10).

